# Recovery after stroke: not so proportional after all?

**DOI:** 10.1101/306514

**Authors:** Thomas M.H. Hope, Karl Friston, Cathy J. Price, Alex P. Leff, Pia Rotshtein, Howard Bowman

## Abstract

**Background:** The proportional recovery rule asserts that most stroke survivors recover a fixed proportion of lost function. Reports that the rule can be used to predict recovery, extraordinarily accurately, are rapidly accumulating. Here, we show that the rule may not be as powerful as it seems.

**Methods:** We provide a formal analysis of the relationship between baseline scores (X), outcomes (Y) and recovery (Y-X), to highlight the shortcomings of the proportional recovery rule, and illustrate those problems with simulations in which synthetic recovery data are derived from different types of recovery processes.

**Findings:** When the correlation between baseline scores and recovery is stronger than that between baselines scores and outcomes, the former can create an inflated impression of how predictable outcomes really are given baseline scores. This often happens when outcomes are less variable than baseline scores, as is common in empirical studies of recovery after stroke. Moreover, we cannot use the results of these correlations to distinguish proportional recovery from recovery which is either not consistently proportional, or not proportional at all.

**Interpretation:** Analyses relating baseline scores to subsequent change are a minefield: our formal analysis applies as consistently outside the area of stroke as it does within it. One implication of our analysis is that the proportional recovery rule is not as predictive of real recovery after stroke as recent empirical studies suggest. Another is that different analytical methods will be required to ascertain whether recovery is even proportional at all.

## 1. INTRODUCTION

Stroke clinicians and researchers have long known that the severity of patients’ initial symptoms after stroke is related to their long term outcomes^1^. Recent studies have suggested that this relationship is stronger than previously thought: that most patients recover a fixed proportion of lost function during the first few months after stroke. Evidence in support of this ‘proportional recovery rule’ is rapidly accumulating, as recently summarised by Stinear^2^. In five separate studies since 2015^3–7^, researchers used the Fugl-Meyer test to assess stroke patients’ upper limb motor function within two weeks after stroke onset, and then again either three or six months post-stroke. The results were consistent with earlier observations^8,9^ that most patients recovered around 70% of lost function. Taken together, these studies report highly consistent recovery in over 500 patients, spread across different countries with different approaches to rehabilitation, regardless of the patients’ ages at stroke onset, stroke type, sex, or therapy dose^2^. And there is increasing evidence that the rule also captures recovery from post-stroke impairments of lower limb function^10^, attention^11,12^, and language^12,13^, and may even apply generally across cognitive domains^14^. Even rats appear to recover proportionally after stroke^15^.

These results are striking because of the extraordinary accuracy with which the proportional recovery rule appears to predict patients’ recoveries. Most of these studies report that the rule explains more (and sometimes much more) than 80% of the variance in empirical recovery: when predicting behavioural performance in humans, these effect sizes are unprecedented. In 2015, Winters and colleagues^3^ reported that recovery predicted using the rule could explain a staggering 95% of the variance in the recovery of 146 stroke patients. Like many of its counterparts^7,10,12^, this study also reported a group of (65) ‘non-fitters’, who did not make the predicted recovery. But if non-fitters can be distinguished at the acute stage, as the authors’ results also suggest, the implication is that we can predict most patients’ recovery near-perfectly, given baseline score alone. Stroke researchers are used to thinking of recovery as a complex, multi-factorial process^16,17^. If the proportional recovery rule is really as powerful as it seems, much of what we thought we knew about recovery from stroke will need to be revisited^2,11,18^.

In what follows, we argue that the proportional recovery rule is not as powerful as it seems, because current analyses in this domain are all potentially confounded by mathematical coupling.

## 2. MATHEMATICAL COUPLING

Mathematical coupling^19,20^ refers to situations in which the same variable is employed to calculate both the independent and the dependent variables in an analysis. In analyses relating baselines to change, change is defined as ‘outcome scores minus baseline scores’: baseline scores are both the independent variable, and a component of the dependent variable. For the sake of brevity in what follows, we define ‘baselines’ = X, ‘outcomes’ = Y, and ‘change’ (or recovery) = A: i.e. delta, or outcomes minus baselines. The ‘correlation between baselines and outcomes’ is r(X, Y), and the ‘correlation between baselines and change’ is r(X, Δ). Finally, we define the ‘variability ratio’ as the ratio of the standard deviation (σ) of Y to the standard deviation of X (σ_y_/σ_x_). X and Y are construed as lists of scores, with each entry being the performance of a single patient at the specified time point. We assume that performance improves as the numbers increase, so r(X, Δ) will be negative if recovery is proportional to lost function (because those who lose the most have smaller X and larger A). One can equally formulate the relationship with ‘lost function’ (maximum X minus actual X) instead of ‘baseline score’, but while this makes r(X, Δ) positive if recovery is proportional to lost function, it is otherwise entirely equivalent.

It has long been known that, when X and Y are independent, random variables drawn from the standard normal distribution, r(X, Δ) ≈ −0.71^21^. In this canonical example of mathematical coupling, there is a strong correlation between baselines and change, even though baselines and outcomes are entirely unrelated: r(X, Y) ≈ 0.

Another expression of this example, which is more germane to our current concerns, is that in this case, r(X, Δ) makes Y appear more predictable than it really is. A strong correlation between two variables typically implies that we can use each to predict the other: in this case, that we can use X to predict Δ. In the limit as our sample grows, out-of-sample predicted Δ (pΔ) should tend toward the least-squares line defined by r(X, Δ), so the correlation between pΔ and empirical Δ should have the same magnitude as r(X, Δ) (see Supplementary Appendix A, proposition 8, bullet point 1). Correlations between predicted and empirical data are a popular measure of predictive accuracy: the stronger the correlation, the better the predictions. Since change is precisely the difference between baselines and outcomes, a result suggesting we can predict change accurately naturally encourages the expectation that we can predict outcomes *equally* accurately, by adding ‘known baselines’ to ‘accurately predicted change’ to produce ‘accurately predicted outcomes’. The expectation is that the correlation between these predicted outcomes and empirical outcomes will be as strong as the correlation between predicted and empirical change, or more formally: |r(X + pΔ, Y)| ≈ |r(X, Δ)|.

However, as might be obvious from our canonical example of mathematical coupling, this expectation is a fallacy: in fact, | r(X + pΔ, Y)| = | r(X, Y)|. No matter how accurately we can seemingly predict recovery, the correlation between outcomes calculated using those predictions, and empirical outcomes, has the same strength as the correlation between baselines and outcomes (see Supplementary Appendix A, proposition 8, bullet 2). In our canonical example, the impression that we can predict change relatively well (because r(pΔ, Δ) ≈ 0.71), is spurious because we cannot use those predictions to predict outcomes even nearly as well (because r(X, Y) ≈ 0).

## 3. MATHEMATICAL COUPLING AND MEASUREMENT NOISE

Krakauer and Marshall recently argued that our canonical example of mathematical coupling is not a proper null hypothesis for empirical analyses of recovery after stroke^18^. This is because the canonical example employs independent baselines and outcomes, whereas empirical baselines and outcomes are dependent; repeated measures from the same patients. Using True Score theory^22^, these authors advance a more plausible alternative, in which ‘true’ baselines become ‘true’ outcomes via some ‘true’ recovery trajectory, and observed baselines and outcomes are just the true states plus measurement noise. From this perspective, the null hypothesis is that baselines are linked to outcomes via random recovery trajectories, and under this hypothesis, it is argued that r(X, Δ) ≈ 0 regardless of the magnitude of recovery: i.e. the spurious effects described previously do not emerge. Only with extreme measurement noise, so the argument goes, will dependent measures behave like independent measures, with the consequent emergence of spurious r(X, Δ). Since the most popular measurement tool in empirical studies of proportional recovery, the Fugl-Meyer scale, is reliable^23^, it follows that those empirical results are probably not spurious^18^.

One weakness in this argument is that it needs to be made again for every new assessment tool employed to measure proportional recovery. For example, neither the subset of the Western Aphasia Battery^24^ used to measure proportional recovery from post-stroke language impairments^13^, nor the letter cancellation task^25^ used to measure proportional recovery from visuospatial neglect^11^, are as reliable as the Fugl-Meyer scale. But whether or not those arguments can be made, we contend that there are many more – and more likely – situations in which r(X, Δ) may be misleadingly strong, regardless of measurement noise.

## 4. MATHEMATICAL COUPLING REGARDLESS OF MEASUREMENT NOISE

For any baselines (X) and outcomes (Y), it can be shown^19^ that:

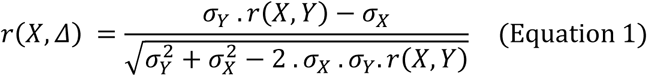

A formal proof of Equation 1 is provided in Supplementary Appendix A (see, proposition 4 and theorem 1); its core consequence is that r(X, Δ) is a (somewhat complex) function of r(X, Y) and σ_y_/σ_x_. To illustrate that function, we ran a series of simulations in which r(X, Y) and σ_y_/σ_x_ were varied independently: a Matlab script for this is in Supplementary Appendix B, and the results are illustrated in Figure 1.

**Figure 1:**
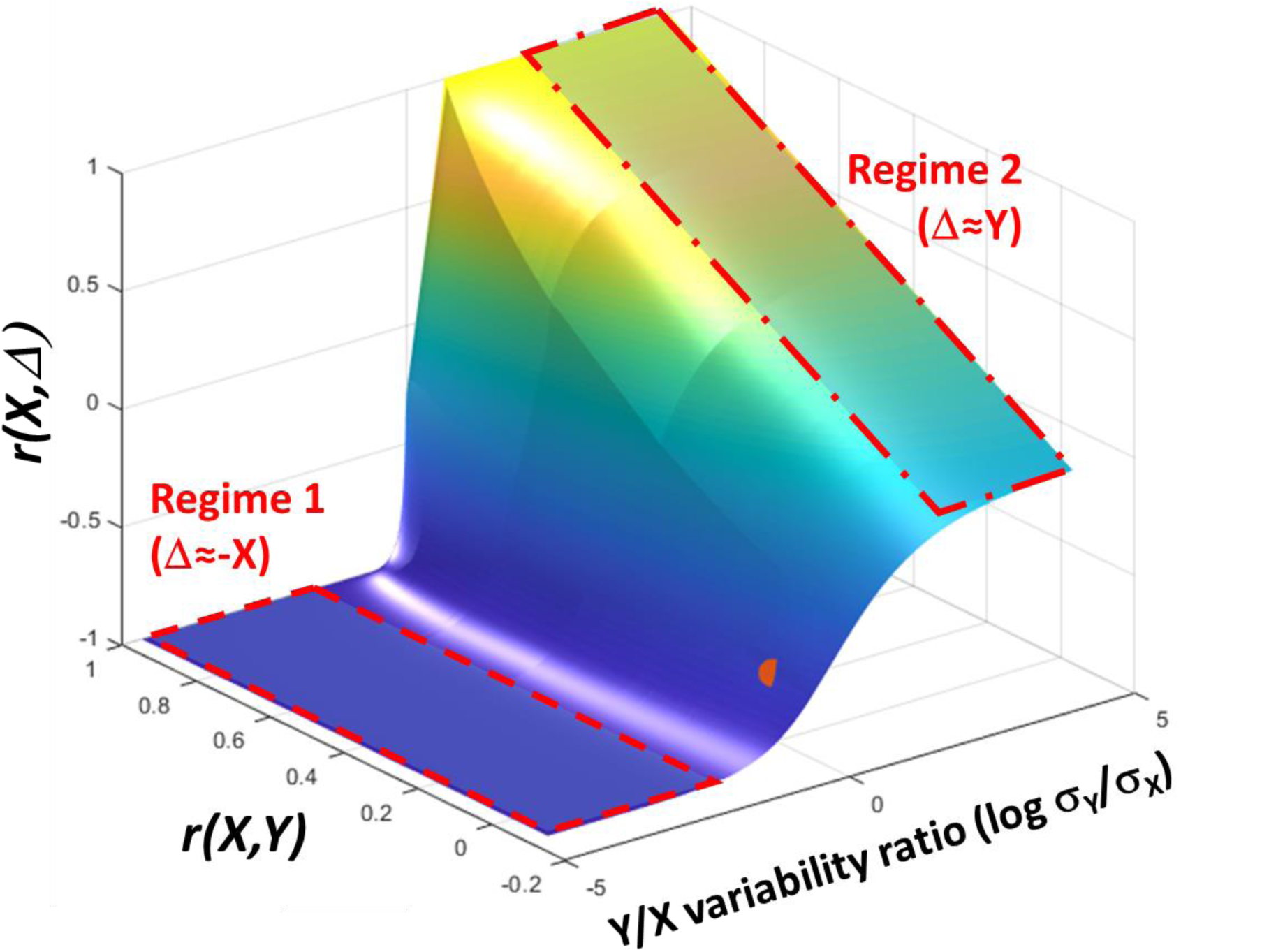
The relationship between r(X, Y), r(X, Δ) and the variability ratio of Y to X. Note that the x-axis is log-transformed to ensure symmetry around 1 (i.e. when X and Y are equally variable, and log(σ_y_/σ_x_) = 0). The two major regimes of Equation 1 are also marked in red. In Regime 1, the imprint of baselines (X) dominates, and r(X, Δ) tends toward r(X, −X) (i.e. −1). In Regime 2, the correlations of interest are more tightly related, and r(X, Δ) ≈ r(X, Y). And there is a transition between the two regimes when the variability ratio is not dramatically skewed one way or the other – though even this transition allows for misleadingly strong r(X, Δ), as evidenced by the fact that our canonical example of mathematical coupling occurs in this region. This is indicated by the red mark, which corresponds to the situation where X and Y are independent random normal variables: i.e. r(X, Y) ≈ 0; log variability ratio = 0; and r(X, Δ) ≈ −0.71. Proposition 7 in Supplementary Appendix A provides a justification for unambiguously using a ratio of standard deviations in this figure, rather than σ_y_ and σ_x_ as separate axes.

**Table 1.**

| REGIME | VARIABILITY OF Y ($\sigma_Y$) | VARIABILITY OF X ($\sigma_X$) | $\Delta [= Y-X]$ | $r(X, \Delta) [=r(X, Y-X)]$ |
| --- | --- | --- | --- | --- |
| 1 | Smaller | Larger | $Y-X \approx -X$ | $r(X, Y-X) \approx r(X, -X) = -1$ |
| 2 | Larger | Smaller | $Y-X \approx Y$ | $r(X, Y-X) \approx r(X, Y)$ |

Our canonical example of mathematical coupling (i.e., when r(X, Y) = 0, σ_y_/σ_x_ = 1 and r(X, Δ) ≈ −0.71) is consistent with equation 1, and corresponds to the red sphere marked in figure 1. One way to understand this surface more generally, is to consider what different variability ratios do to the correlation between X and Y-X (i.e. Δ): see Table 1. When Y is more variable than X, Y contributes more variance to Y-X and r(X, Y-X) tends toward r(X, Y): this is Regime 2. In contrast, when Y is less variable than X, X contributes more variance to Y-X, and r(X, Y-X) tends toward r(X, −X), the autocorrelation between X and itself: this is Regime 1 (also see proposition 9 in Supplementary Appendix A). As an illustration of Regime 1, consider a set of synthetic baselines and outcomes connected by proportional recovery. With no measurement noise, r(X, Y) = 1 and r(X, Δ) = −1 (Figure 2a); but with shuffled outcomes data, r(X, Y) ≈ 0, but r(X, Δ) = −0.96 (Figure 2b). Indeed, r(X, Δ) will be extreme in any data where σ_y_ is significantly smaller than σ_x_.

**Figure 2:**
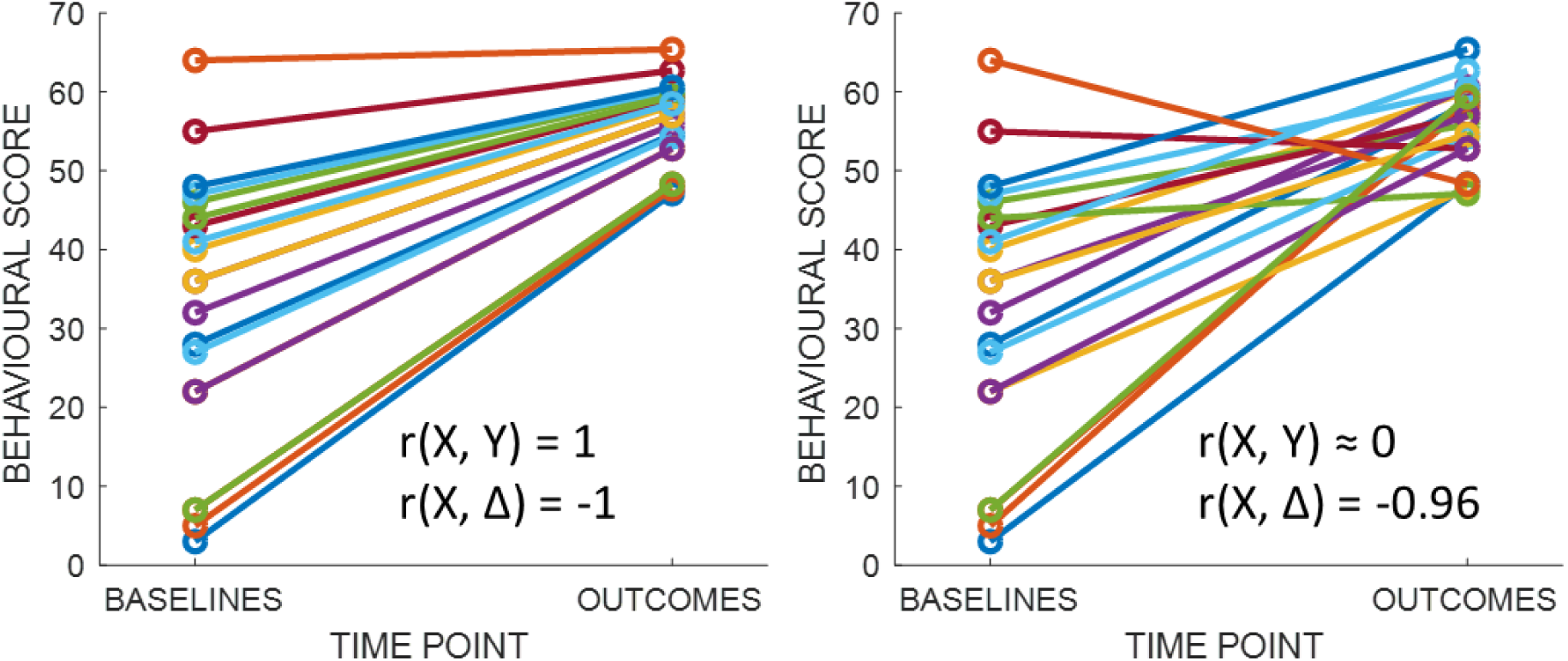
Illustrating how | r(X, Δ)| > | r(X, Y)| when X is more variable than Y. In both panels, X is a vector of 20 numbers drawn from a random uniform distribution in the range 1-65. In the left panel, Y is calculated using the proportional recovery rule: i.e. Y = X + ((66-X) × 0.7). In this case, the correlation between X and Y is 1 and the correlation between X and change is −1. In the right panel, Y is randomly shuffled: X and Y are now uncorrelated, but the correlation between X and change remains extremely strong (r = −0.96).

## 5. RE-EXAMINING STUDIES OF PROPORTIONAL RECOVERY AFTER STROKE

These relationships between r(X, Y), r(X, Δ) and σ_Y_/σ_X_ merit a re-examination of the proportional recovery rule for stroke. The only study in this literature that reports individual patients’ behavioural data is that by Zarahn and colleagues^9^, who reported recovery from hemiparesis in 30 stroke patients. When measured across the whole sample, r(X, Y) = 0.80 and r(X, Δ) = −0.49; this changes to r(X, Y) = 0.75 and r(X, Δ) = −0.95, after removing the 7 patients (23%) who did not fit the rule. Calculating predicted recovery (pΔ) from the least-squares line for r(X, Δ), we get r(pΔ, Δ) = 0.95, but r(X + pΔ, Y) = 0.75. By removing those 7 non-fitters, Zarahn and colleagues dramatically increased the apparent predictability of recovery, while actually reducing the apparent predictability of outcomes – and reducing the variability ratio, from 0.88 in the full sample to 0.36 in the reduced sample of fitters. Notably, the residuals or errors for both correlations (r(X, Y) and r(X, Δ)) are identical (see Figure 3), and in fact this is always true (see Supplementary Appendix A, proposition 10). Since r(X, Y) and r(X, Δ) assign different effect sizes to the same errors, r(X, Δ) is misleadingly strong.

**Figure 3:**
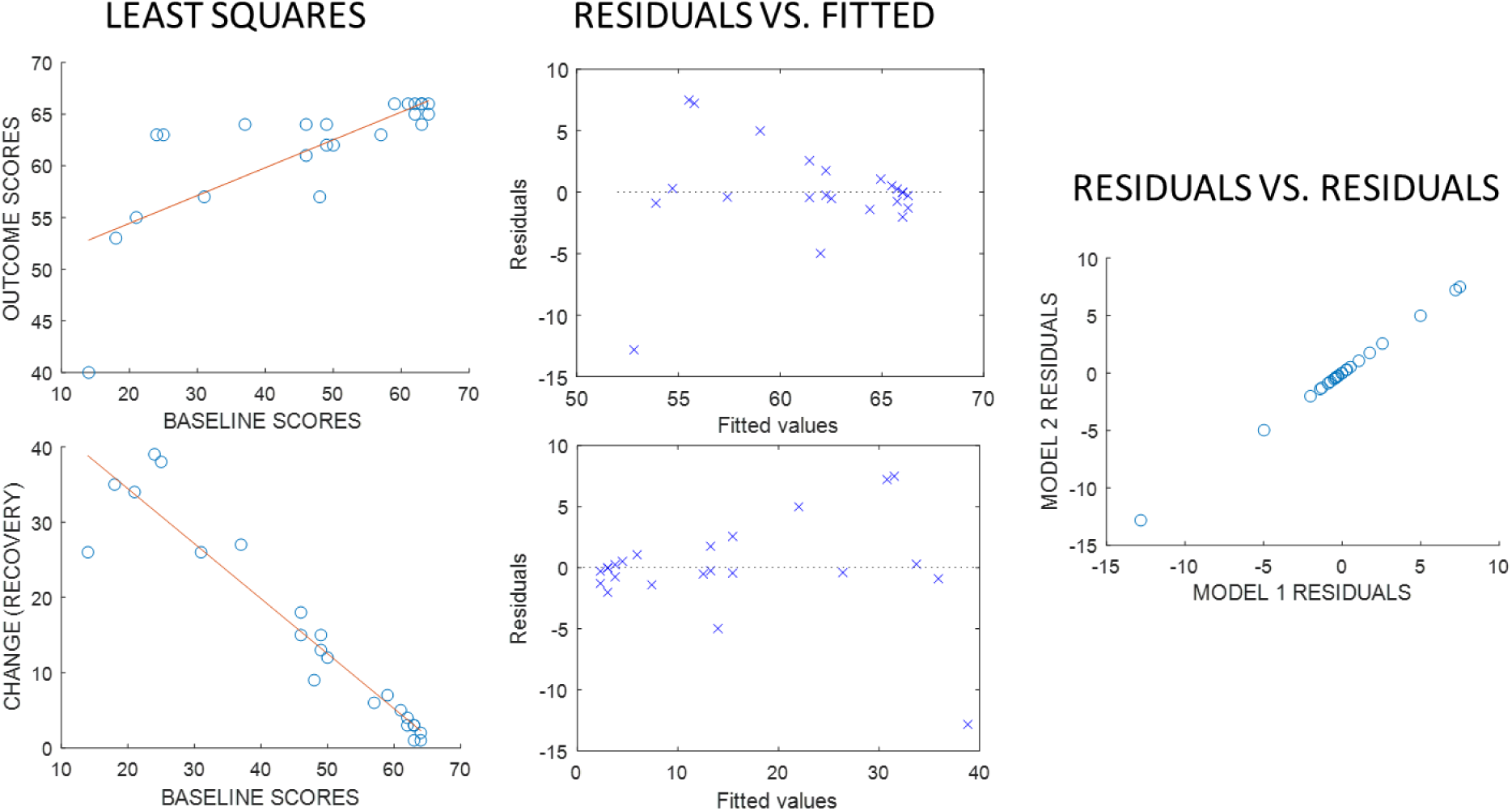
(Left) Least squares linear fits for analyses relating baselines to (upper) outcomes and (lower) recovery, using the fitters’ data reported by Zarahn and colleagues^9^. (Middle) Plots of residuals relative to each least squares line, against the fitted values in each case. (Right) A scatter plot of the residuals from the model relating baselines to change, against the residuals from the model relating baselines to outcomes: the two sets of residuals are the same.

We can also use equation 1 to reinterpret studies which do not report individual patient data. One example is the first study to apply the proportional recovery rule to aphasia after stroke^13^. Here, baseline scores explained 81% of the variance in recovery (r(X, Δ) ≈ −0.9), and the variability ratio was 3.7 (outcomes) / 7.7 (baselines) = 0.48. By equation 1, r(X, Y) was at most 0.78 here, but could also have been zero; without a report of r(X, Y), we cannot know how predictable Y really was, given X. Similar concerns emerge from the recent measurement of proportional recovery in 182 rats (from a total sample of 593)^15^. Here, the variability ratio was 3.2 (outcomes) / 4 (baseline) = 0.8, baseline scores explained 49.9% of the variance in recovery (r(X, Δ) ≈ −0.71). By equation 1, this implies that r(X, Y) was either much stronger (>0.95) or considerably weaker (~0.29) than their r(X, Δ). Here again, we have learned less about the predictability of outcomes than the headline effect size suggests.

Some studies report inter-quartile ranges (IQR) instead of standard deviations for their data; accepting some room for error, we can still estimate variability ratios from those IQRs. The study by Winters and colleagues^3^ is an example, which reports proportional recovery in 146 patients (from a total sample of 211). Here, r(X, Δ) = −0.97 and σ_y_/σ_x_ = 6 (follow-up) / 38 (baseline) = 0.158. By equation 1, this small ratio means that r(X, Δ) will be at least as extreme as that reported regardless of the r(X, Y). And the same issues arise in the recent study of proportional recovery from lower-limb impairments, by Veerbeck and colleagues^26^. Here, σ_y_/σ_x_ = 0.438 and r(X, Δ) ≈ −0.88; equation 1 implies that r(X, Δ) could be at least that strong regardless of r(X, Y). The headline effect sizes in these studies do not tell us how predictable outcomes really were, given baseline scores.

Finally, while r(X, Δ) can be misleading if it is extreme relative to r(X, Y), the reverse is also true. For example, the only study in this literature to employ outcomes as the dependent variable, rather than recovery, reports that r(X, Y) ≈ 0.8 and σ_y_/σ_x_ = 23.4 (follow-up) /19.5 (baseline) = 1.2 in their ‘combined’ group of 76 patients. By equation 1, r(X, Δ) = −0.05 in this case. These authors only claimed that a subset of their sample recovered proportionally, but their full sample seems quite consistent with the null hypothesis, of random recovery, implied by a True Score analysis of recovery data^18^.

Notably, most of the studies referenced in this section reported outcomes scores which were less variable than baseline scores. This is the predicted pattern if recovery is proportional to lost function, because in this case those who lose the most will also recover the most, but this scenario also encourages extreme r(X, Δ) regardless of r(X, Y) (i.e. Regime 1 from Figure 1). Even if patients really do recover proportionally, r(X, Δ) can encourage an inflated impression of how well the rule predicts their recovery. We found potentially misleading results in all of the studies that reported enough information for us to make the judgement. The most common reason why we could not make a judgement was when baselines were only related to recovery through multivariable models^4,11,12^. But inflation in one variable’s effect size will also inflate the multivariable model’s effect size: we cannot estimate the magnitude of that inflation confidently for any particular multivariable model, but we can predict that it’s also there in those models.

As a consequence, we suggest that future studies of proportional recovery should at least report both r(X, Y) and r(X, Δ). However, even when both are reported, we contend that analyses like this cannot convincingly demonstrate that recovery is proportional: this is the focus of the next section.

## 6. HOW DO PATIENTS REALLY RECOVER?

To illustrate why r(X, Y) and r(X, Δ) cannot convincingly identify proportional recovery, we ran a series of simulations in which these correlations were calculated from synthetic data, generated by different types of recovery process. All of the simulations assume that baselines and outcomes are dependent, and that each is observed with some measurement noise, and all also assume our samples are large (N = 1,000); the distortions induced by small samples are well documented elsewhere^27^, and will not concern us here.

Our simulations vary: (a) the magnitude of measurement noise; (b) the real variation in participants’ recovery trajectories (i.e. ‘recovery noise’); (c) the presence or absence of boundaries on the measurement scale (i.e. ceiling and floor effects); and (d) whether or not ‘non-fitters’ (whose ‘recovery’ is less than expected) are removed from the analysis. Three different types of recovery are considered: (i) random recovery; (ii) constant recovery (i.e. patients recover a fixed amount); and (iii) proportional recovery. Results (r(X, Y) and r(X, Δ)) are mean values from 100 repetitions of the same simulations: the Matlab script required to run them is in Supplementary Appendix C.

Figure 4 displays the results: r(X, Y) and r(X, Δ) with increasing recovery noise, with each configuration of recovery mechanism and other constraints. As expected given our discussion in section 3, when measurements are noisy (Figure 4, row 1), mathematical coupling dominates and all of the recovery regimes yield similar results. With more reliable measurements (Figure 3, row 2), the results are more promising, in that one can reasonably hope to use these correlations to distinguish proportional recovery from other types of recovery. However, these simulations make the assumption that the scale used to measure performance is infinitely long. In practice, these scales are always finite, and the Fugl-Meyer scale in particular is known to exert ceiling effects on patients with mild-to-moderate hemiparesis^23^. When we include ceiling effects (Figure 4, row 3), the results derived from constant recovery are similar to those derived from proportional recovery. These similarities persist when we also remove non-fitters prior to calculating the correlations (Figure 4, last row), which acts to preserve strong r(X, Δ) despite significant recovery noise. Indeed, in our final simulation of proportional recovery (row 4, column 2), r(X, Δ) ≈ −0.94 even when the standard deviation of the recovery noise is >0.35. Since the mean proportion is 70% here (0.7), the 95% confidence intervals (mean ± two standard deviations) on the proportion of lost function actually recovered include both 0% and 100%.

**Figure 4:**
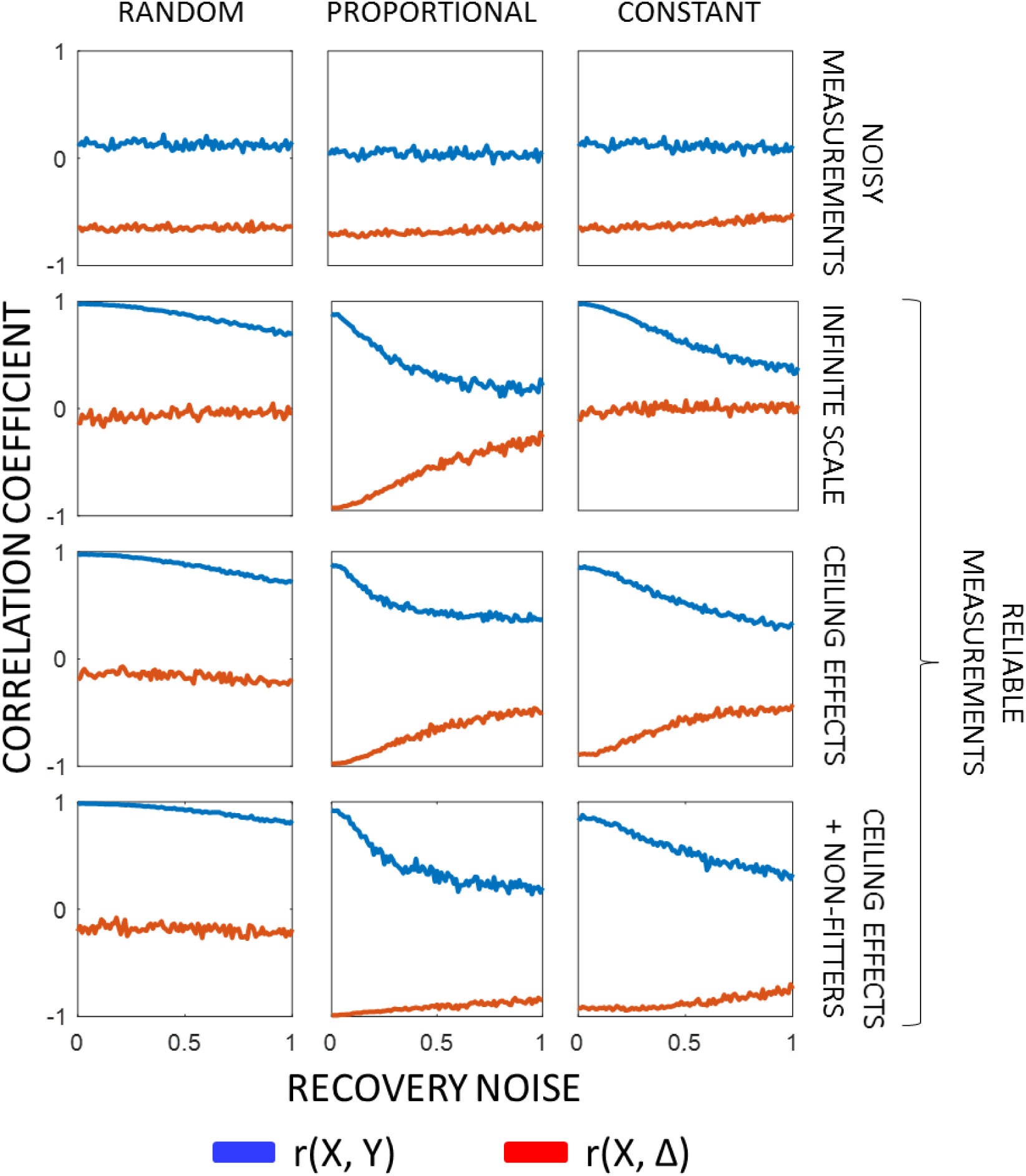
Simulation results. Plotting r(X, Y) and r(X, Δ) as a function of increasing true variability in recovery trajectories, where recovery is: (a) random; (b) proportional; or (c) constant. The top row displays results when the variance of measurement noise is large relative to the variance of baselines. The second row presents results when measurement noise is minimal. The third row presents results when measurement noise is minimal but there are ceiling effects on the measurement scale. And the last row presents results when measurement noise is minimal, ceiling effects are applied, and ‘non-fitters’ are removed: i.e. the ~19% (this is the figure in ^3^) of the sample whose actual recovery deviates most strongly and negatively from the least squares line defined by r(X, Δ).

These results demonstrate that, with more realistic constraints on our generative models of empirical data, r(X, Y) and r(X, Δ) cannot distinguish proportional recovery from recovery which is either not consistently proportional, or is not proportional at all.

## 7. Discussion

The proportional recovery rule is attractive partly because it formalises the popular, clinical intuition that initial symptom severity is by far the most, and perhaps the only, important predictor of recovery after stroke. The rule is also striking because it implies that recovery is simple and consistent across patients (non-fitters notwithstanding), and because that implication appears to be justified by strong empirical results^2^. That incredible, empirical power has even begun to encourage the search for some deeper, general recovery mechanism which cuts across cognitive domains^18^. But our results suggest that the rule may not be as powerful as it seems.

We are not claiming that the rule is wrong. Our analysis suggests that the empirical results to date do not confirm that it holds or how well, but those results are still consistent with the rule’s predictions. The rule is also falsifiable: a recent study of recovery during the first week after stroke onset found no evidence of proportional recovery^28^. Nevertheless, we contend that estimates of the rule’s predictive power – those headline effect sizes for r(X, Δ) or r(pΔ, Δ) – deliver a misleadingly optimistic impression of how predictable outcomes really are. And a misleadingly optimistic impression here encourages misleadingly pessimistic assessments elsewhere: if baseline scores really do predict many patients’ recoveries near-perfectly, therapeutic interventions will appear to be redundant for those patients.

Much of the work in this area has focused on the distinction between fitters and non-fitters to the proportional recovery rule; distinguishing the groups via analyses of the residuals for r(X, Δ). This is a circular definition, but it defines groups which can then be distinguished in non-circular ways^5^. Since the residuals for r(X, Y) and r(X, Δ) are always the same (see Supplementary Appendix A), it follows that our concerns about effect sizes have no bearing on the validity of work to distinguish fitters from non-fitters. Nevertheless, extreme r(X, Δ) for fitters will naturally encourage the assumption that we can at least predict those fitters’ outcomes near-perfectly. Our analysis suggests that this assumption is wrong.

In summary, our results suggest that recovery from stroke might not be as proportional as it seems. Strong correlations between baseline scores (or predicted change) and subsequent change can drive a misleadingly optimistic impression of how predictable outcomes really are. This situation appears to be common, both in our simulations and in the empirical literature on the proportional recovery rule. Moreover, these analyses cannot distinguish proportional recovery from recovery which is either not consistently proportional, or not proportional at all. Quite how to make this distinction with confidence, remains an open question: this is a subject for future work.

## 8. A PARELLEL CHALLENGE?

As this manuscript approached completion, we discovered that another researcher, Robinson Kundert, had distributed a ‘pre-paper’, describing a parallel challenge to the proportional recovery rule, at a meeting. None of the authors attended that meeting, but we learned of it through colleagues who did attend. Our understanding is that this challenge is focused on the consistency (or lack thereof) of the proportion of lost function that patients recover. In this sense, that work may be complementary to our focus on predictive power. We look forward to studying this challenge after it is published.

## ACKNOWLEDGMENTS

This study was supported for the Medical Research Council (MR/M023672/1, MR/K022563/1), the Wellcome Trust (091593/Z/10/Z), and the Stroke Association (TSA PDF 2017/02). The funders had no participation in the design and results of this study.

## Supplementary Appendix A: formal relationships between the correlations

We present a very simple and general formulation of the proportional recovery concept. We assume two variables *X’* and *Y’* corresponding to performance at initial test (*X’*) and at second test (*Y’*). These will be represented as column vectors, with each entry being the performance of a single patient and vector lengths being *N ∈ ℕ*. Performance improves as numbers get bigger, up to a maximum, denoted *Max*, which corresponds to no discernible deficit. Severity is measured as difference from maximum, i.e. *Max – X’*.

The two variables (*X’* and *Y’*) could be specialised to more detailed formulations: e.g., true score theory or with an explicit modelling of measurement or state error. However, this would not impact any of the derivations or inferences that follow. Indeed, the results that we present would hold even in the complete absence of measurement noise, which has been considered the main concern for the validity of quantifications of proportional recovery.

### Demeaning

Without loss of generality, we work with demeaned variables. That is, where over-lining denotes mean, we define new variables as,

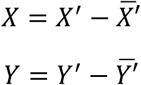

This also means that recovery, i.e. *Y − X*, will be demeaned, since, using proposition 1, the following holds.

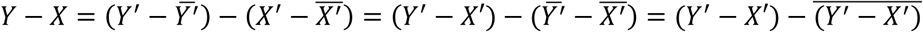

#### Proposition 1

Let *V* and *W* be vectors of the same length, denoted *N*. Then, the following holds,

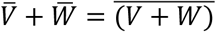

with 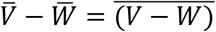 as a trivial consequence.

#### Proof

By distributivity of multiplication through addition and associativity of addition, the following holds.

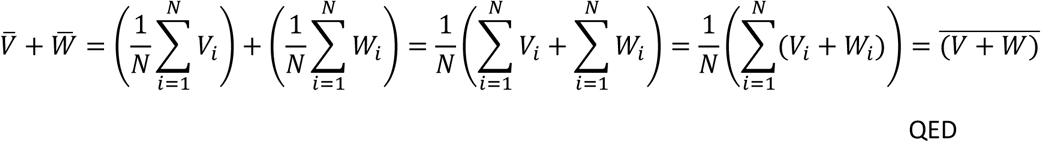

### Correlations

There are two basic correlations we are interested in, (1) the correlation between initial performance and performance at second test, i.e. *r*(*X, Y*), and (2) the correlation between initial performance and recovery, i.e. *r*(*X, Y − X*) = *r*(*X*, Δ). The latter of these is the key relationship, and we would expect this to be a negative correlation; that is, as initial performance is smaller (i.e. further from *Max*), the larger is recovery. (One could also formulate the correlation as *r*((*Max − X*), *Y − X*), which would flip the correlation to positive, but the two approaches are equivalent).

Our main correlations are defined as follows,

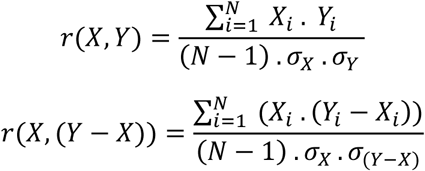

### Standard Deviation of a Difference

We need a straightforward result on the standard deviation of a difference.

#### Proposition 2

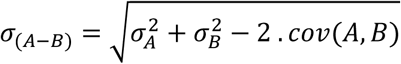

#### Proof

The result is a direct consequence of the following standard result from probability theory, e.g. see Ross, S. M. (2014). *Introduction to probability and statistics for engineers and scientists*. Academic Press.,

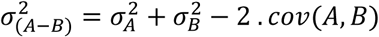

### Key Results

The following proposition enables us to express the key correlation, *r*(*X*, (*Y − X*)), in terms of covariance of its constituent variables.

#### Proposition 3

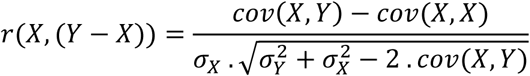

#### Proof

Using distributivity of multiplication through addition, associativity of addition, the definition of covariance and proposition 2, we can reason as follows.

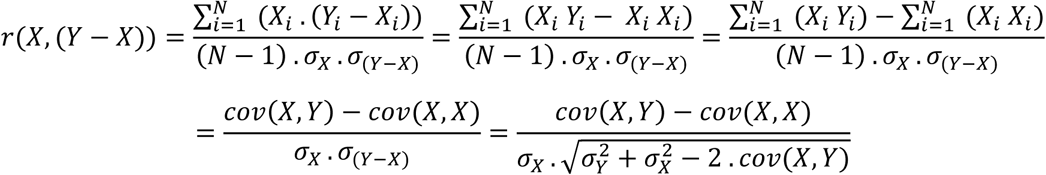

QED

It is straightforward to adapt proposition 3 to be fully in terms of correlations.

#### Proposition 4

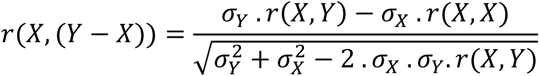

#### Proof

Straightforward from proposition 3 and definition of correlations, which gives the relationship

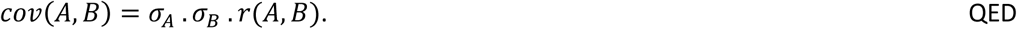

Scale Invariance

The next set of propositions justifies working with a standardised *X* variable.

#### Lemma 1

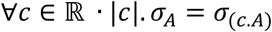

#### Proof

Using distributivity of a multiplicative constant through averaging, 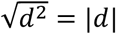 and distributivity of square root through multiplication, we can reason as follows.

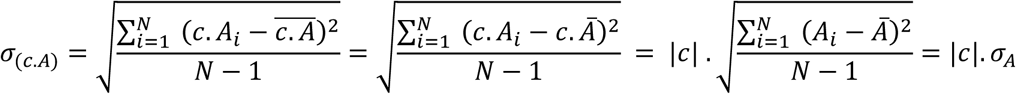

QED

#### Proposition 5 (Invariance to scaling)

The absolute magnitude of a correlation is not changed by scaling either variable by a constant, i.e.

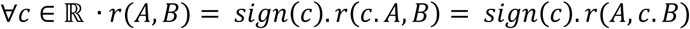
where *sign*(*d*) = *if* (*d* < 0) *then* − 1 *else* + 1.

#### Proof

For any *c* ∈ ℝ, using distributivity of multiplication through mean and addition, and lemma 1, the following holds,

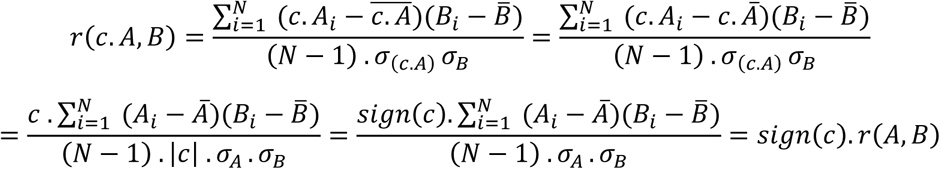

Then, one can multiply both sides by *sign*(*c*) to obtain *r*(*A, B*) = *sign*(*c*)*. r*(*c. A, B*). Additionally, as correlations are symmetric, *sign*(*c*)*. r*(*c. B, A*) = *sign*(*c*)*. r*(*A, c. B*), and the full result follows.

QED

#### Corollary 1

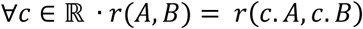

#### Proof

Follows from twice applying proposition 5, and that *sign*(*c*)^2^ *=* +1. QED

#### Proposition 6

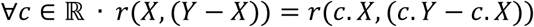

#### Proof

We can use distributivity of multiplication through subtraction and corollary 1 to give us the following.

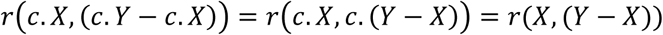

QED

It follows from proposition 6 that we can work with a standardised *X* variable, since,

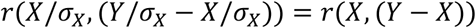

#### Proposition 7 (Sufficiency of variability ratio)

Assume two pairs of variables: *X*_1_*, Y*_1_ and *X*_2_*, Y*_2_, such that, *r*(*X*_1_, *Y*_1_) = *r*(*X*_2_*, Y*_2_), then,

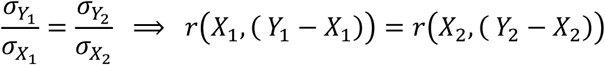

#### Proof

The proof has two parts.

1) We consider the implications of equality of ratio of standard deviations. Firstly, we note that,

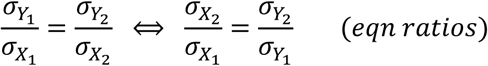

Secondly, using eqn ratios, we can argue as follows,

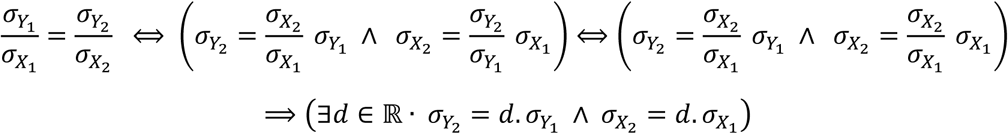

2) Using 4, the fact that *r*(*X*_1_*, Y*_1_) = *r*(*X*_2_*, Y*_2_), the property just derived in part 1), with 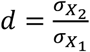 and rules of square roots, we can reason as follows,

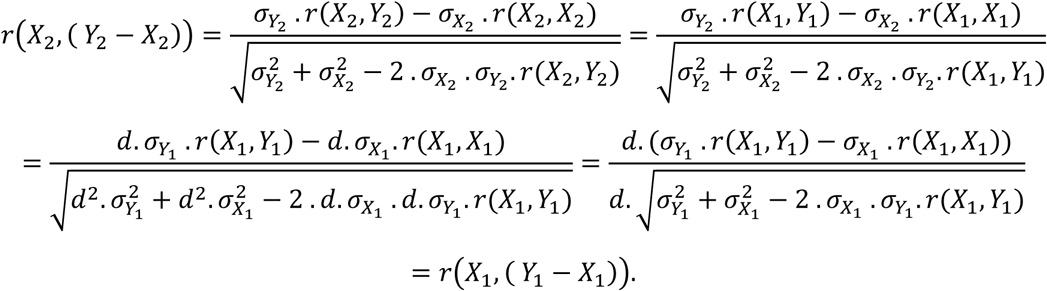

QED

#### Proposition 8

If Δ = *Y − X* and *p*Δ=*X. β*, where *β∈*ℝ, then,

1) *r*(*p*Δ,Δ)= *sign*(*β*). *r*(*X*, Δ); and

2) *r*(*X* + *p*Δ,Y) = *sign*(*1 + β*)*.r*(*X*,Y).

#### Proof

Both results are easy consequences of proposition 5.

1) *r*(*p*Δ, Δ) = *r*(*X. β*, Δ) = *sign*(*β*)*. r*(*X*, Δ).

2) *r*(*X + p*Δ,Y) = *r*((*X +* (*X.β*)),Y) = *r*((*X*. (1 + *β*)),Y) = *sign*(1 *+ β*)*.r*(*X*,Y) = *r*(*X*,Y).

QED

### Main Findings

#### Theorem 1

Since *X* will be standardised, we can adapt the finding in proposition 4, to give us the key relationship we need,

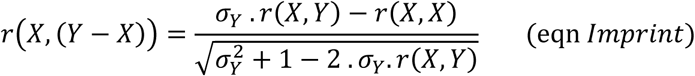

#### Proof

Immediate from proposition 4. QED

Theorem 1 shows clearly that *r*(*X*, (*Y − X*)) is fully defined by the correlations *r*(*X, Y*) and *r*(*X, X*), along with the variability of *Y*. The correlation of *X* with itself, i.e. *r*(*X, X*), is a prominent aspect of this equation, which drives its oddities. *r*(*X,X*) reflects the coupling in the equation that arises because *X* appears in both the terms being correlated in *r*(*X*, (*Y − X*))*. r*(*X, X*) is of course a constant, i.e. 1 for any *X*, so in fact, *σ_Y_* and *r*(*X, Y*), are the only variables; accordingly, their size determines the extent to which the imprint of *X* in *Y − X* drives *r*(*X*, (*Y − X*)).

This leads to the key observation that, as *σ_Y_* gets smaller, *r*(*X*, (*Y − X*)) tends towards *−r*(*X, X*), which equals −1. In other words, as the variability of Y decreases, the imprint of *X* becomes increasingly prominent. This is shown in the next proposition.

#### Proposition 9

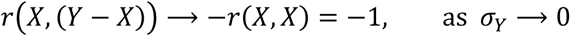

#### Proof

The right hand side of equation *Imprint*, has five constituent terms, two in the numerator and three in the denominator. Of these five, three are products with the standard deviation of *Y*, i.e. *σ_Y_*. Assuming all else is constant, as *σ_Y_* reduces, the absolute value of each of these three terms reduces towards zero. The rate of reduction is different amongst the three, but they will all decrease. Accordingly, as *σ_Y_* decreases, *r*(*X*, (*Y − X*)’) becomes increasingly determined by the two terms not involving *σ_Y_*, and thus, it tends towards 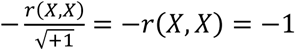.

QED

### Equality of Residuals

An important finding of section 5 of the main text, is that the residuals resulting from regressing Y onto X are the same as regressing Y-X onto X. We show in this section, that this equality of residuals is necessarily the case.

We focus on the following two equations,

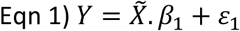

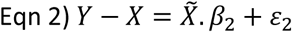

where 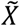 is the *N* × 2 matrix, with first column being *X* and second being the *N* × 1 vector of ones (which provides the intercept term); *β*_1_ and *β*_2_ are 2×1 vectors of parameters and *Y*, *X, ε*_1_ and *ε*_2_ are *N* × 1 vectors. As in the rest of this document, *Y* and *X* are our (demeaned) initial and outcome variables, while *ε*_1_ and *ε*_2_ are our residual error terms.

#### Proposition 10

If we assume that *β*_1_ and *β*_2_ are fit with ordinary least squares, with *ε*_1_ and *ε*_2_ the associated residuals, then, *ε*_1_ *− ε*_2_.

#### Proof

Under ordinary least squares, the parameters are set as follows.

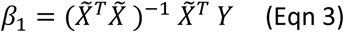

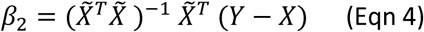

We start with the second of these, and using left distributivity of matrices, and then substituting Eqn 3, we obtain the following.

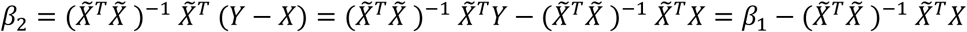

Using the fact that the variable *X* is demeaned, we can now evaluate the main term here as follows,

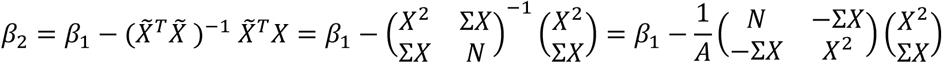

where *X*^2^ is the dot product of *X* with itself, Σ*X* is the sum of the vector *X*, and *A = NX*^2^ − Σ*X*Σ*X* is the determinant of the matrix being inverted. From here we can derive the following,

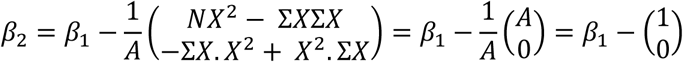

We can then substitute this equality for *β*_2_ in eqn 2 and re-arrange to obtain,

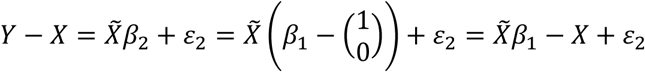

It follows straightforwardly from here that,

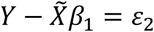

i.e. *ε*_1_ *− ε*_2_, as required.

QED

Proposition 10 shows that the residuals resulting from fitting equations 1 and 2 will be the same. A consequence of this is that the error variability will be the same. As a result of this, the factor that determines whether more variance is explained when regressing *Y* onto *X* or when regressing *Y − X* onto *X*, is the variance available to explain. That is, the relative variance of *Y* and *Y − X* drive the *R*^2^ values of these two regressions. This then implicates the variance of *Y* and *X* and in fact their covariance (which impacts the variance of *Y − X*).

More precisely, we can state the following.

1) If 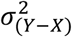 is big relative to 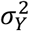, then regressing *Y − X* onto *X* will explain more variability than regressing *Y* onto *X*.

2) If 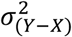 is small relative to 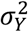, then regressing *Y − X* onto *X* will explain less variability than regressing *Y* onto *X*.

## Supplementary Appendix B: illustrating the relationship between the correlations

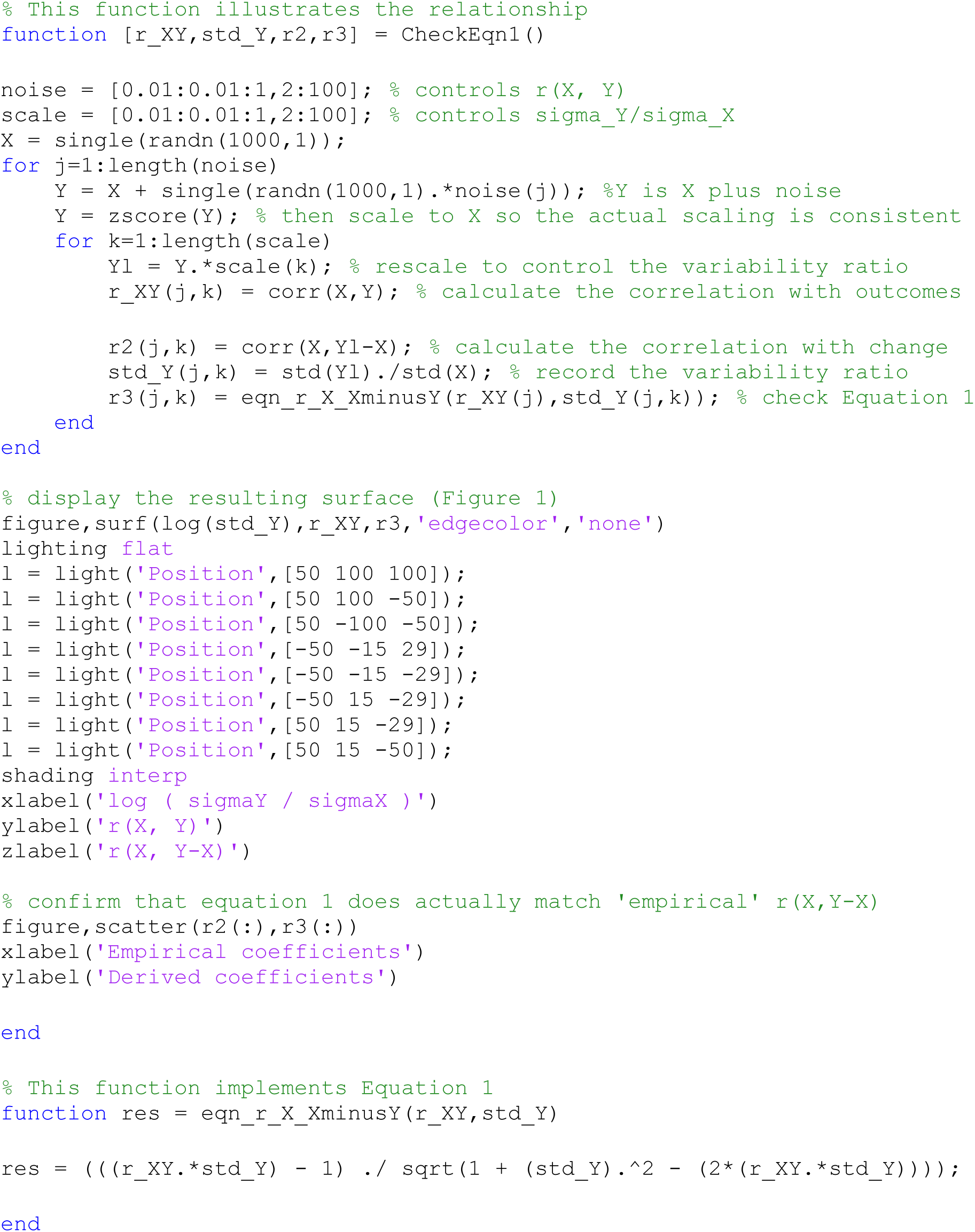

## Supplementary Appendix C: simulating recovery

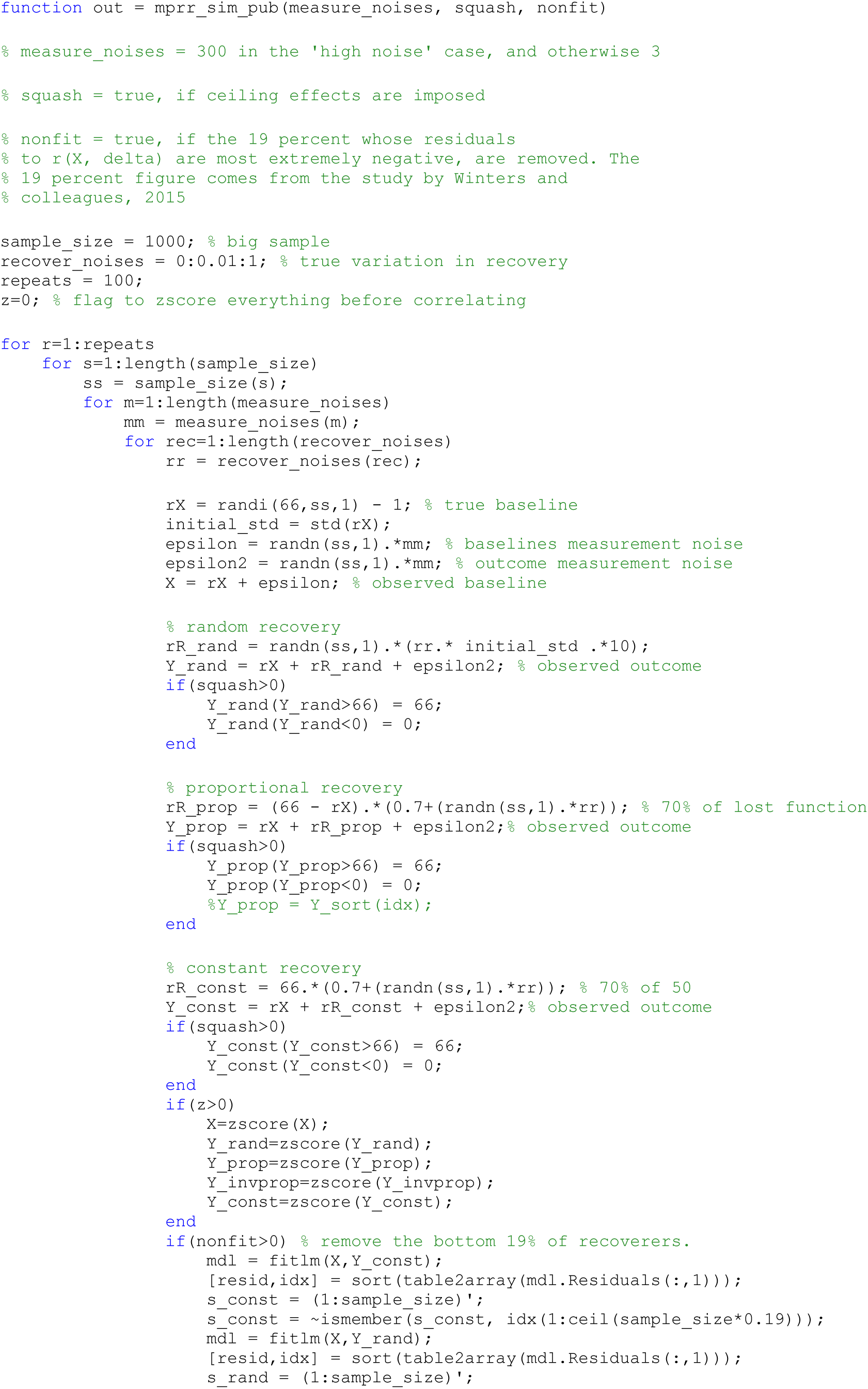

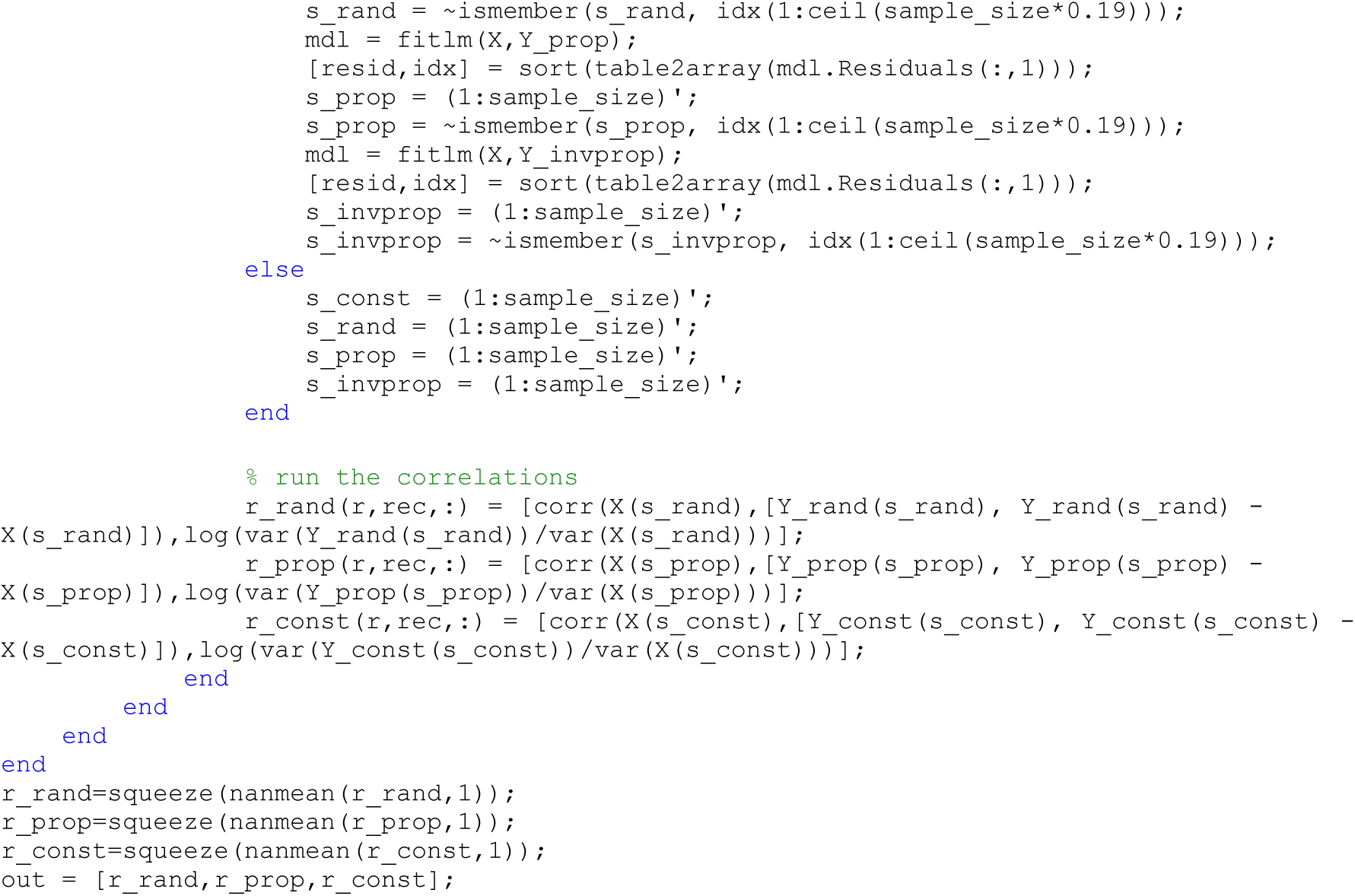

